# Sex-biased transcriptome in embryonic mouse cortices under *Pax6* haploinsufficiency highlights Pbdc1 as a candidate regulator

**DOI:** 10.1101/2025.10.19.683266

**Authors:** Shyu Manabe, Shohei Ochi, Takako Kikkawa, Sharmin Naher, Mai Saeki, Kohdai Yamada, Hidetaka Kosako, Yusuke Kishi, Tatsuya Sawasaki, Noriko Osumi

## Abstract

Mutations in *Pax6*, encoding a transcription factor essential for brain patterning and neurogenesis, have been linked to female-biased cortical malformations and behavioral abnormalities, yet the molecular basis remains unclear. Here we show that *Pax6* haploinsufficiency (*Sey*/+) produces more pronounced sex-biased alterations in the transcriptomes and cytoarchitecture of embryonic mouse cortices than those in wild-type and homozygous mutants (*Sey*/*Sey*). *Pbdc1*, a previously uncharacterized X-linked gene implicated in autism, is selectively upregulated in *Sey*/+ females and proximity-dependent protein-protein interaction analysis reveals Pbdc1 interacts with RNA-splicing factors. Moreover, *Pbdc1* overexpression reduces intermediate progenitor cells in the developing cortex. ChIP-qPCR further demonstrates Pax6 and BAF occupancy at the *Pbdc1* promoter in WT embryos of both sexes and CUT&Tag shows H3K4me3 elevation selectively in *Sey*/+ females. Our findings indicate that partial loss of Pax6 shapes the embryonic cortical transcriptomes and cytoarchitecture in a sex-dependent manner and highlight *Pbdc1* as a candidate regulator of sex-biased corticogenesis.

## Introduction

The mammalian brain exhibits region-specific sexual dimorphism. A long-standing model posits that a “default” female-type brain is masculinized by a perinatal testosterone surge, establishing biases for sex-specific behaviors (e.g., mounting in males and lordosis in females) that emerge around and after puberty^1^. More recently, this purely hormonal framework has been challenged by evidence that embryonic gene programs can prime sexual differentiation before birth; for example, hypothalamic Ptf1a expression is required for brain sexual differentiation^2^. However, whether comparable genetic programs operate in other regions, such as the cerebral cortex, remains unclear.

Sex differences are also prominent across neurodevelopmental disorders (NDDs)^3^. Beyond Tourette syndrome^4^ and attention-deficit/hyperactivity disorder (ADHD)^5^, autism spectrum disorder (ASD) shows a marked male bias in diagnosis^6^. Clinical profiles also diverge by sex: females with ASD more often exhibit internalizing symptoms (e.g., anxiety and depression), whereas males more commonly display externalizing features (e.g., hyperactivity and repetitive/restricted behaviors)^7,8^. Nevertheless, many ASD model studies either focus on males or pool sexes, leaving the molecular basis of sex differences in ASD prevalence and phenotype poorly defined.

Among ASD-associated genes curated by Simons Foundation Autism Research Initiative (SFARI)^9^ is *PAX6*, which encodes a transcription factor highly expressed in the embryonic cortex and essential for neural patterning and corticogenesis^10–15^. Deletion of 11p13 region, which include *PAX6* gene, is associated with WAGR (Wilms tumor, aniridia, genital ridge defects, mental retardation) syndrome^16^, a subset of whom also meet diagnostic criteria for ASD^17^. Additional studies have also linked *PAX6* variants to ASD^18,19^. In rodents, homozygous *Pax6* mutations are perinatally lethal, whereas heterozygotes show ASD-relevant behavioral endophenotypes: impaired social behaviors in *Pax6* mutant rats (*rSey*^2^*/+*)^20^ and altered ultrasonic vocalizations and hyperactivity in *Pax6* mutant mice (*Sey/+*)^21^.

Notably, a few *Pax6*-related phenotypes are reported to be sex-biased. Magnetic resonance imaging (MRI) has revealed greater cortical volume reduction in *rSey*^2^/+ females relative to wild-type (WT) than in males^22^, and ultrasonic vocalizations were more reduced in *rSey*^2^/+ females^20^. In mice, *Sey/+* females exhibit more pronounced abnormalities in the anterior commissure morphology^23^. These observations suggest sex-dependent effects by partial reduction of Pax6 on cortical development; however, it remains unresolved whether sex differences are already embedded within embryonic cortical programs, the developmental window when Pax6 is pivotal.

Here, we address this gap by profiling sex-biased transcriptional programs in the embryonic cortex of *Sey*/*+* mice. Using RNA-seq with orthogonal validation, we find that sex differences in the transcriptomes are most pronounced in *Sey/+* relative to WT and homozygous mutants (*Sey*/*Sey*). We identify *Pbdc1*, a previously uncharacterized X-linked gene associated with ASD, as selectively upregulated in *Sey*/+ females and in both sexes of *Sey*/*Sey*, with strong expression in neural stem/progenitor cells (NSPCs) and intermediate progenitor cells (IPs). Proximity- dependent protein-protein interaction (PPI) analysis implicates Pbdc1 in mRNA- splicing/processing pathways, and manipulating *Pbdc1* levels in the developing cortex perturbs IPs: knockdown increases IP population, whereas overexpression decreases it without altering NSPC population. Furthermore, we show robust occupancy of Pax6 and BAF chromatin remodeling factors at the promoter region of *Pbdc1* in WT males and females, accompanied by a female-specific increase of H3K4me3, an epigenetic mark linked to transcriptional activation, in *Sey*/+. Together, these findings indicate that *Pax6* haploinsufficiency elicits sex-specific transcriptional changes during corticogenesis and highlight Pbdc1 as a candidate mediator of females-biased cortical vulnerability in *Pax6* mutant embryos.

## Results

### *Pax6* haploinsufficiency amplified sex-biased transcription in the E14.5 telencephalon

To examine sex differences in gene expression, we performed bulk RNA-seq on the telencephalon from WT, *Sey*/+ and *Sey*/*Sey* mouse embryos at embryonic day 14.5 (E14.5). We focused on E14.5 because sex-biased transcriptomes in the murine telencephalon are more pronounced at this stage relative to E11.5^24^. UMAP of sample-level transcriptomes showed that *Sey*/*Sey* males and females exhibited similar expression profiles. Although one sample each from WT male, WT female, *Sey*/+ male and *Sey*/+ female clustered apart from their representative groups, most of WT males and females intermingled, whereas *Sey*/+ males and females segregated into two distinct clusters (**Fig. 1a**). Consistently, the number of sex-differentially expressed genes (sex-DEGs; *FDR* < 0.1) was greatest in *Sey*/+ (**Fig. 1b**). Qualitative comparison of sex-DEGs across genotypes revealed that WT and *Sey*/*Sey* exhibited comparable sex differences; 80% of the sex-DEGs were shared between WT and *Sey*/*Sey*, and these genes were predominantly X- or Y-chromosome linked (**Fig. 1c**). In contrast, *Sey*/+ harbored a unique set of sex-DEGs that were not detected in WT or *Sey*/*Sey*.

**Fig. 1:**
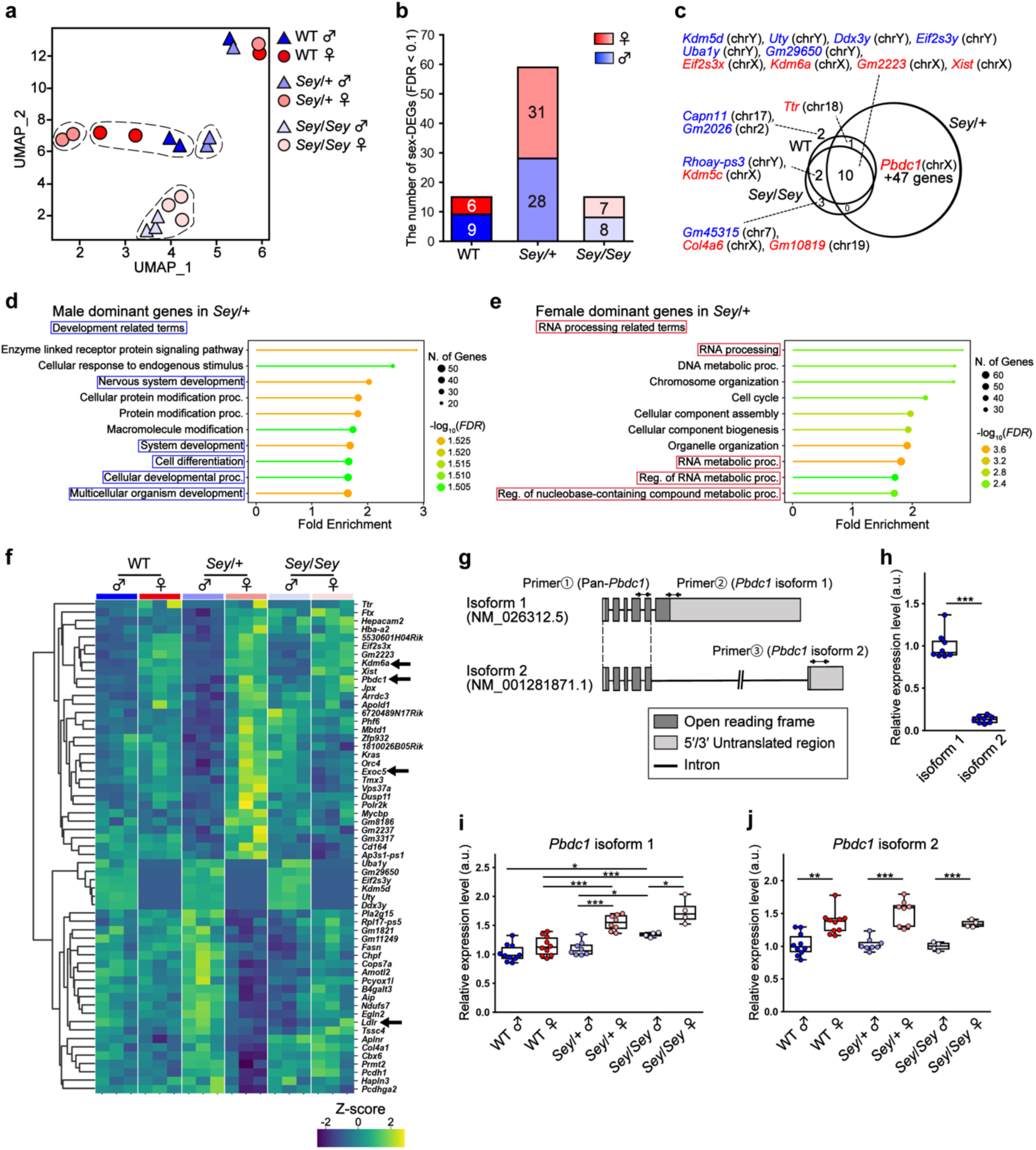
Sex differences in the gene expression patterns of the *Pax6* mutant telencephalons. **a** UMAP plots of the RNA-seq data from male and female telencephalons of WT, *Sey*/+ and *Sey*/*Sey*. **b** The number of sex-DEGs in each genotype (*FDR* < 0.1). **c** Venn diagram showing the overlap of sex-DEGs among the three genotypes. Blue, highly expressed in males; red, highly expressed in females. **d, e** GO (Biological Process) analyses of male-dominant (**d**) and female-dominant (**e**) DEGs in *Sey*/+ embryos (*FDR* < 0.25). **f** Heatmap showing normalized (z-score) expression levels of 59 sex-DEGs between males and females in *Sey*/+ embryos. Arrows indicate genes that have been previously reported to be associated with ASD. **g** Schematic illustration of the RT-qPCR primers designed to detect pan-Pbdc1 (primer ①), isoforms 1 (primer ②) and isoform 2 (primer ③) of murine *Pbdc1*. **h** RT-qPCR analysis of *Pbdc1* isoform1 and isoform2 mRNA expression levels in the telencephalons of WT male embryos. **i, j** RT-qPCR analysis of *Pbdc1* isoform 1 (**i**) and isoform 2 (**j**) mRNA expression levels in the telencephalons of WT, *Sey*/+ and *Sey*/*Sey* embryos. \**p* < 0.05, \*\**p* < 0.01, \*\*\**p* < 0.001; determined by two-tailed Student’s *t*-test followed by Benjamini-Hochberg multiple testing correction. The line in the middle of box plots represent median, the lower and upper bounds of boxes indicate the first and third quartiles, and end of whiskers denote the minimum and maximum values.

Given this pronounced sex bias in *Sey*/+, we focused subsequent analyses on this genotype. Gene Ontology (GO) analyses for Biological Process indicated that genes upregulated in *Sey*/+ males (i.e., female-down) were enriched for development-related terms (e.g.f, “Nervous system development”, “System development” and “Cell differentiation”; **Fig. 1d**). Conversely, genes upregulated in *Sey*/+ females (i.e., male-down) were enriched for RNA-processing-related terms (e.g., “RNA processing”, “RNA metabolic process” and “Regulation of RNA metabolic process”; **Fig. 1e**).

To refine candidate drivers of the *Sey*/+ sex differences, we visualized normalized expression levels (z-score) of the 59 sex-DEGs in *Sey*/+ across all the samples as a heatmap and intersected them with previously reported ASD-related gene sets (**Fig. 1f**). Four genes overlapped; three female-dominant sex-DEGs in *Sey*/+, i.e., *Kdm6a*^25^, *Pbdc1*^26^ and *Exoc5*^27^, and one male-dominant sex-DEG in *Sey*/+, i.e., *Ldlr*^28^. Reverse transcription quantitative PCR (RT-qPCR) corroborated female-biased expression of *Kdm6a* and *Pbdc1* across WT, *Sey*/+ and *Sey*/*Sey*, whereas *Exoc5* and *Ldlr* did not show consistent sex differences (**Supplementary Fig. 1a**).

We next quantified isoform-specific expression of *Kdm6a* and *Pbdc1* in our bulk RNA-seq dataset using *Salmon*^29^. Two predominantly expressed *Kdm6a* isoforms (ENSMUST00000044484/NM_001310444.2 and ENSMUST00000225336 [no RefSeq match]), showed similar female-biased expression across genotypes (**Supplementary Fig. 1b**). In contrast, *Pbdc1* ENSMUST00000033577 (NM_026312.5, hereafter isoform 1) exhibited the largest sex difference specifically in *Sey*/+ compared to WT and *Sey*/*Sey,* whereas ENSMUST00000119477 (NM_001281871.1, isoform 2) showed smallest sex difference in *Sey*/+ (**Supplementary Fig. 1d**). Isoform specific RT-qPCR confirmed that *Pbdc1* isoform 1 was expressed at substantially higher levels than *Pbdc1* isoform 2 (**Fig. 1g, h**). While isoform 2 maintained a female-biased pattern across genotypes, *Pbdc1* isoform 1 was markedly upregulated in *Sey*/+ females and elevated in both sexes of *Sey*/*Sey* relative to WT (**Fig. 1i, j**). Collectively, these results indicate that *Pbdc1*, particularly isoform 1, shows sex- and genotype-dependent upregulation, with a pronounced *Sey*/+ female-specific increase, motivating our focus on this molecule in the subsequent analyses.

### *Pbdc1* isoform 1 is enriched in neural stem/progenitor cells

Prompted by a prior report of ∼1.7 Mb deletions on Xq encompassing six genes, including *PBDC1,* in three ASD cases^26^ and the limited knowledge of *Pbdc1* in mammalian corticogenesis, we first characterized spatial and cell-type-specific expression patterns of this gene in the developing mouse cortex. Re-analysis of a public single-cell RNA-seq (scRNA-seq) dataset from E14.5 mouse cortex^30^ revealed robust pan-*Pbdc1* expression in NSPCs and IPs (**Fig. 2a**). A developmental time-course dataset spanning E12-E15^31^ further showed that pan-*Pbdc1* expression peaks in NSPCs during early corticogenesis (E12-14) and declines as neurogenesis progresses (**Fig. 2b**). To validate these patterns in tissue, we performed *in situ* hybridization (ISH) with a pan-*Pbdc1* probe (recognizing both isoforms) on WT female brains. Consistently, signals were strongest in the progenitor-enriched ventricular zone (VZ) and subventricular zones (SVZ) at E11.5 and E14.5, diminishing by E17.5 (**Fig. 2c**). ISH across WT, *Sey*/+ and *Sey*/*Sey* embryos at E14.5 detected pan-*Pbdc1* expression in both males and females, with stronger expression in females across the genotypes, concordant with RT-qPCR for pan-*Pbdc1* (**Fig. 2d**). We next applied RNAscope, a highly sensitive and quantitative fluorescent ISH, specific to *Pbdc1* isoform 1 and detected a significant increase in isoform 1 signal in the cortical VZ of *Sey*/+ females relative to WT at E14.5, in agreement with isoform-specific RT-qPCR for *Pbdc1* isoform 1 (*p* = 0.045, **Fig. 2e, f**). Together, integrative re-analysis of public datasets and *in situ* validations indicate that *Pbdc1* is enriched in NSPCs and IPs, and that isoform 1 exhibits sex- and genotype-dependent upregulation, with a pronounced increase in *Sey*/+ females.

**Fig. 2:**
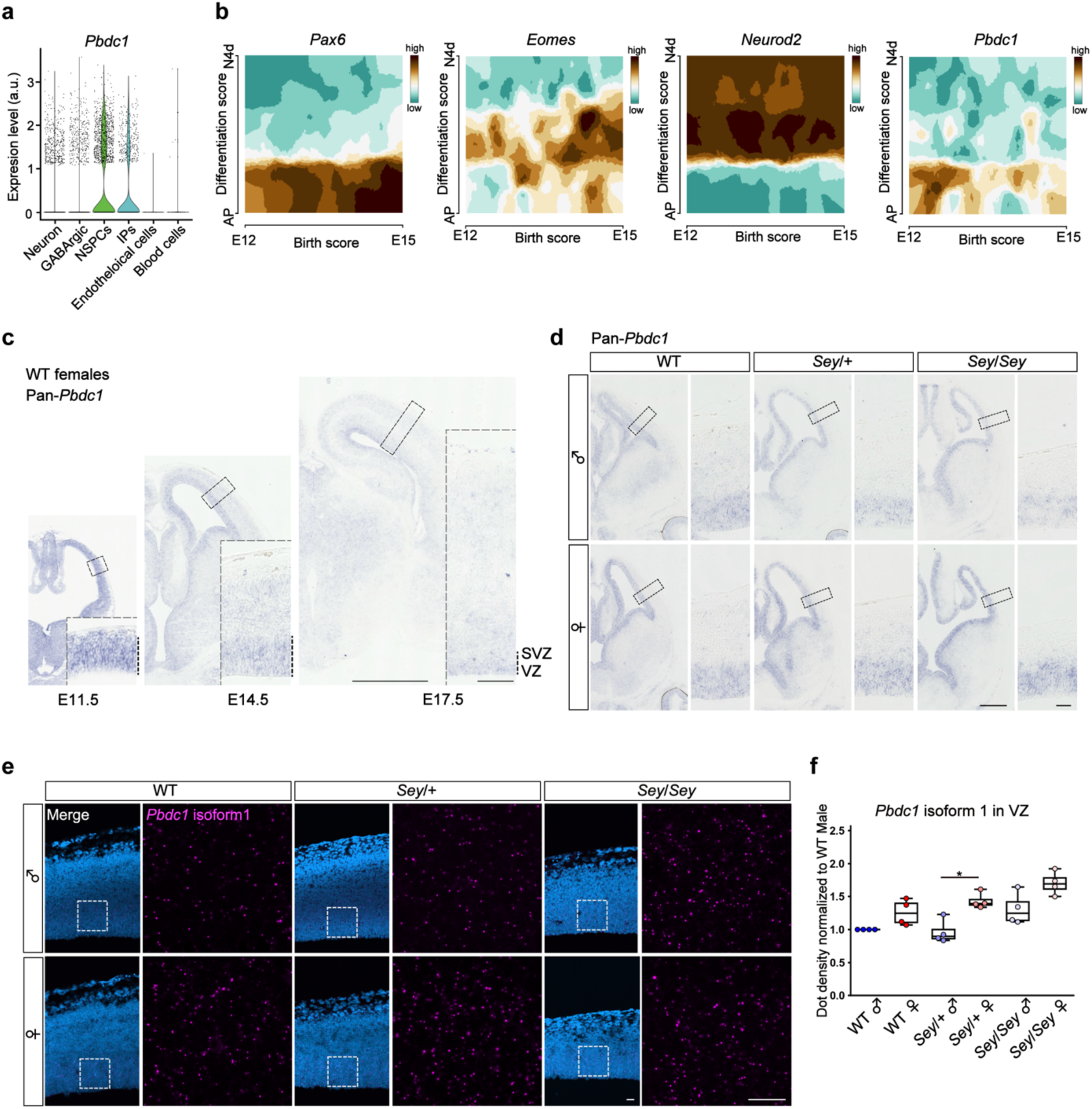
*Pbdc1* expression in the developing cortex. **a** Pan-*Pbdc1* mRNA expression levels in previously published single-cell RNA-seq data from mouse cortex at E14.5^30^. **b** mRNA expression pattern of *Pax6* (NSPCs), *Eomes* (IPs), *Neurod2* (neurons) and pan*-Pbdc1* in mouse cortex from E12 to E15, based on previously published single cell RNA-seq data^31^. **c** *in situ* hybridization of pan-*Pbdc1* mRNA (purple signal) in the telencephalons of WT female embryos at E11.5, E14.5 and E17.5. Boxes denote the zoomed area. Scale bars, lower magnification, 1 mm; higher magnification, 100 µm. **d** *in situ* hybridization of pan-*Pbdc1* mRNA (purple signal) in the telencephalons of WT, *Sey*/+ and *Sey*/*Sey* embryos at E14.5. Scale bars, lower magnification: 500 µm, higher magnification: 50 µm. **e** RNAscope analysis of *Pbdc1* isoform 1 (magenta) in the cortex of WT, *Sey*/+ and *Sey*/*Sey* embryos at E14.5. Nuclei were counterstained with DAPI (cyan). Boxes denote the zoomed area. Scale bars, 25 μm. **f** Quantification of *Pbdc1* isoform 1 dot density in the VZ across sexes and genotype. Dot density was normalized to the value of WT male within the same experimental batch to minimize batch effects. \**p* < 0.05; determined by two-tailed Student’s *t*- test followed by Benjamini-Hochberg multiple testing correction. Abbreviation: NSPCs, neural stem/progenitor cells; AP, apical progenitor cells; IPs, intermediate progenitor cells; N4d, 4- day-old neurons; VZ, ventricular zone; SVZ, subventricular zone. The line in the middle of box plots represent median, the lower and upper bounds of boxes indicate the first and third quartiles, and end of whiskers denote the minimum and maximum values.

### Pbdc1 isoform 1 associates with RNA-splicing machinery

To probe the molecular function of Pbdc1 isoform 1 (hereafter, Pbdc1), we performed proximity-dependent PPI analysis using AirID, a proximity biotinylation enzyme^32^. Mouse neuroblastoma N2a cells stably expressing AGIA-AirID-Pbdc1 or AGIA-AirID-IκBα (validated control bait) were generated by lentivirus transduction. Upon biotin supplementation, proteins in proximity to Pbdc1 or IκBα were biotinylated by AirID and identified by liquid chromatography-tandem mass spectrometry (LC-MS/MS) (**Fig. 3a**). PCA analysis of the biotinylated proteomes showed clear separation of the Pbdc1 and IκBα interactomes, indicating distinct proximal assemblies (**Fig. 3b**). Applying significance and enrichment criteria (*p* < 0.05, Log2(Pbdc1/IκBα) > 2), we refined a high-confidence set of 488 Pbdc1-enriched proteins (**Fig. 3c**). GO analysis revealed strong enrichment for RNA-splicing terms (e.g., “RNA splicing via transesterification reactions” and “RNA splicing via transesterification reactions with bulged adenosine as nucleophile”), as well as RNA processing terms (e.g., “mRNA processing” and “mRNA metabolic process”) (**Fig. 3d, e**). Notably, among 43 Pbdc1-interacting proteins annotated to RNA splicing terms, 29 have reported roles in cell-cycle control or oncogenic progression, including HNRNPK^33^, SNRPA1^34^ and SART1^35^ (**Supplementary Table 1**). Together, these data indicate that Pbdc1, previously uncharacterized in corticogenesis, might associate with RNA-splicing/processing and cell cycle regulating factors, suggesting a role in coupling pre-mRNA-splicing to proliferation/differentiation programs in NSPCs and IPs.

**Fig. 3:**
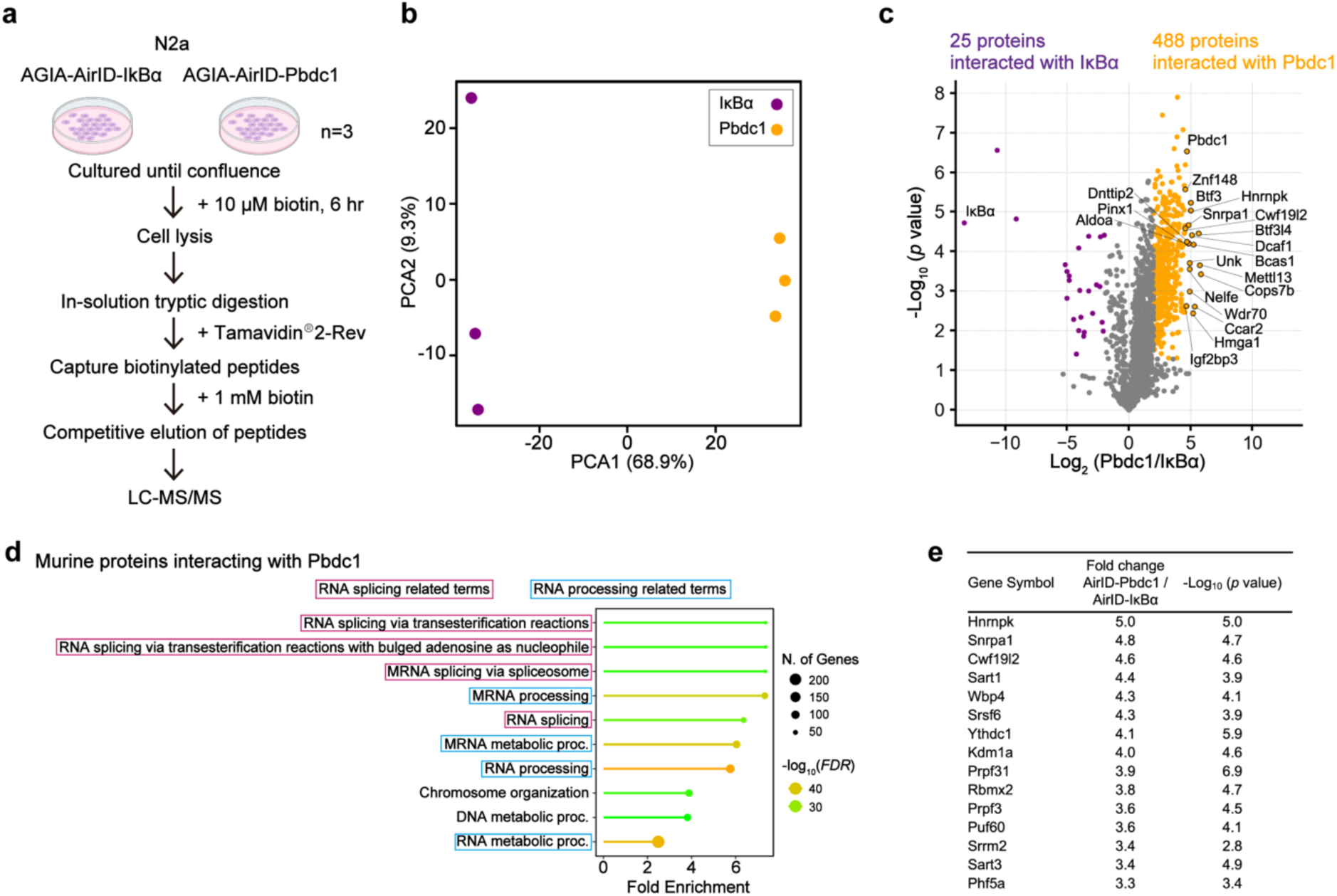
Proximity-dependent protein-protein interaction analysis in AirID-Pbdc1 expressing cells. **a** Schematic illustration for detecting biotinylated proteins using mouse neuroblastoma N2a cells stably expressing AGIA-AirID-Pbdc1 or AGIA-AirID-IκBα. **b** Principal component analysis (PCA) of the proteins which showed interaction with either Pbdc1 or IκBα detected by LC-MS/MS. **c** A volcano plot showing AirID-Pbdc1 vs. AirID-IκBα. The numbers of the significantly enriched proteins (*p* < 0.05; |Log2(Pbdc1/ IκBα)| > 2.0|) are shown. Names of the top 20 proteins enriched in AirID-Pbdc1 and IκBα are indicated. **d** GO (Biological Processes) analysis of proteins enriched in AirID-Pbdc1 (*p* < 0.05; Log2(Pbdc1/IκBα) > 2.0). **e** A list of top 15 proteins increased in AirID-Pbdc1 and involved in RNA-splicing terms.

### Pbdc1 isoform 1 modulates proliferation and differentiation of NSPCs

Given the association of Pbdc1 with RNA-splicing/processing factors linked to cell-cycle control, we asked whether its elevated expression in *Sey*/+ females relates to sex differences in corticogenesis. Cytoarchitectural analyses confirmed genotype-dependent changes in cortical primordium thickness: *Sey*/*Sey* cortices were significantly thinner than WT and *Sey*/+ (WT males vs. *Sey*/*Sey* males, *p* = 0.009; *Sey*/+ males vs. *Sey*/*Sey* males, *p* = 0.005; WT females vs. *Sey*/*Sey* females, *p* = 0.021; *Sey*/+ females vs. *Sey*/*Sey* females, *p* = 0.021; **Fig. 4a-c**), consistent with previous reports^36,37^. No statistically significant male–female differences were detected across genotypes; however, *Sey*/+ females showed a trend toward reduced cortical thickness relative to *Sey*/+ males (*p* = 0.126; **Fig. 4a-c**). Marker analyses further showed that the relative thickness of the Sox2-positive (Sox2^+^) NSPC compartment tended to be higher in *Sey*/+ females than that of *Sey*/+ males (*p* = 0.069) and was significantly increased in *Sey*/*Sey* males compared to *Sey*/+ males (*p* = 6.75×10^-4^), paralleling overall thinning in *Sey*/*Sey* cortices (**Fig. 4d**). In contrast, the number of the Tbr2^+^ IPs was significantly lower in *Sey*/+ females than in *Sey*/+ males (*p* = 0.026) and was reduced in *Sey*/*Sey* of both sexes compared with WT and *Sey*/+ samples, in line with previous observations^38^ (**Fig. 4e**).

**Fig. 4:**
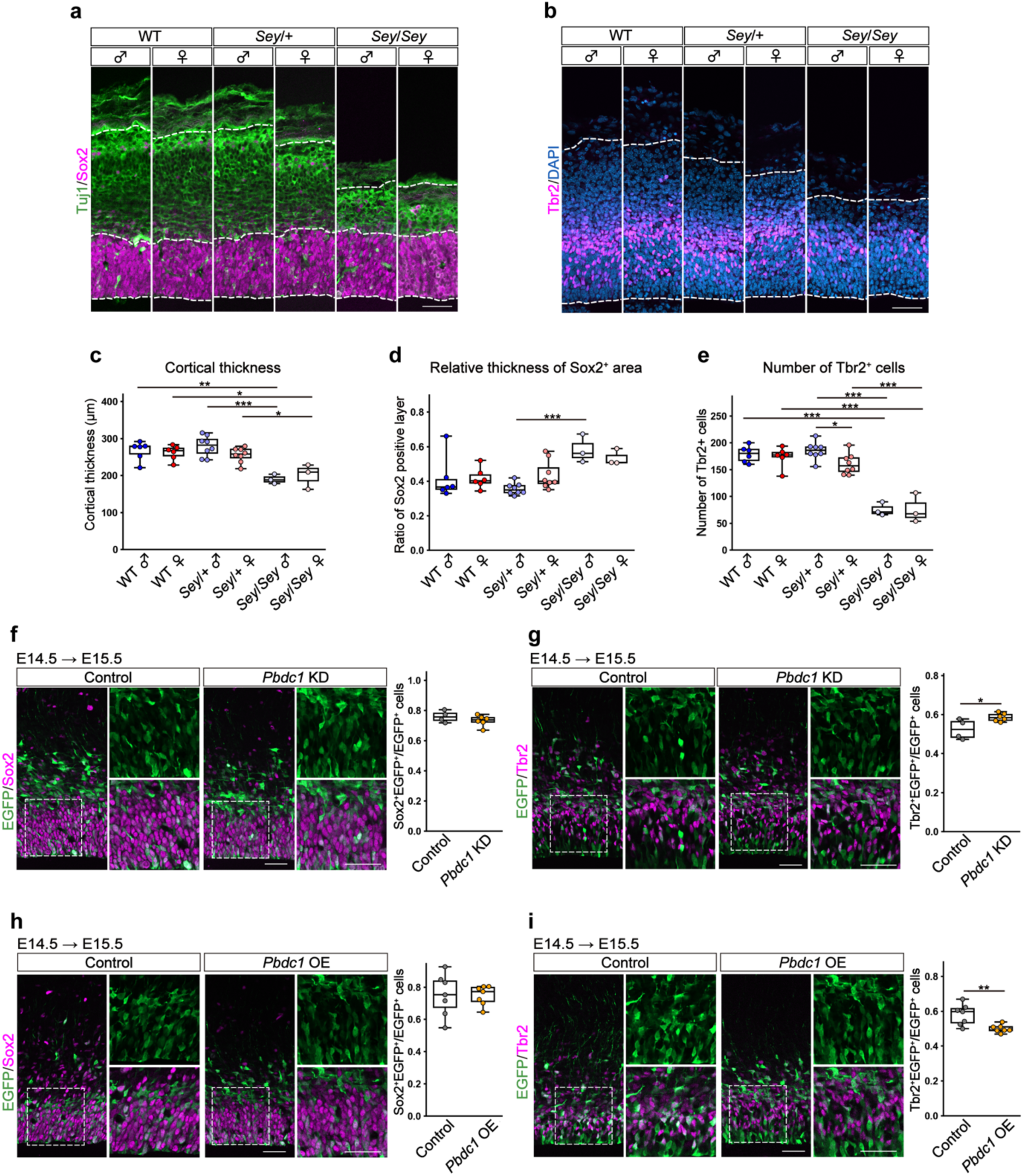
Knockdown and overexpression analyses of Pbdc1 in developing cortex. **a,b** Representative images of cortices stained for Sox2 and Tuj1 (a); Tbr2 and DAPI (b) in WT, *Sey*/+ and *Sey*/*Sey* of both sexes. Scale bars, 50 µm. **c-e** Quantification of cortical thickness (c), Sox2^+^ area relative to total cortical area within the 100 µm wide column (d) and the number of Tbr2^+^ cells within the 100 µm wide column of E14.5 cortices (e). **f, g** Representative images of the control and *Pbdc1* KD cortices stained for Sox2 (f, left) and Tbr2 (g, left), together with EGFP at 24 h after electroporation (E15.5). Boxes of each image denote the zoomed area. Scale bars, 50 µm. Quantification of the relative ratio of Sox2^+^EGFP^+^ cells (f, right) and Tbr2^+^EGFP^+^ cells (g, right) relative to the total EGFP^+^ cells within the 200 μm wide column in E15.5 cortices. **h, i** Representative images of the control and *Pbdc1* OE cortices stained for Sox2 (h, left) and Tbr2 (i, left), together with EGFP at 24 h after electroporation (E15.5). Boxes of each image denote the zoomed area. Scale bars, 50 µm. Quantification of the relative ratio of Sox2^+^EGFP^+^ cells (h, right) and Tbr2^+^EGFP^+^ cells (i, right) relative to the total EGFP^+^ cells within the 200 μm wide column in E15.5 cortices. \**p* < 0.05, \*\**p* < 0.01, \*\*\**p* < 0.001; determined by two-tailed Student’s *t*-test followed by Benjamini-Hochberg multiple testing correction. The line in the middle of box plots represent median, the lower and upper bounds of boxes indicate the first and third quartiles, and end of whiskers denote the minimum and maximum values.

To explore the potential involvement of Pbdc1 in the sex-biased cytoarchitecture, we examined loss- and gain-of-function effects of Pbdc1 on NSPCs and IPs production *in vivo*. First, we employed an *in utero* electroporation (IUE) technique to knockdown (KD) *Pbdc1* by delivering a small interfering RNA (siRNA) targeting exon 6 of murine *Pbdc1* isoform 1, together with an EGFP expression vector, into the mouse neocortex at E14.5. *Pbdc1* mRNA was successfully reduced at E15.5, 24 h post-IUE (**Supplementary Fig. 2a**). Under this condition, the proportion of EGFP^+^Sox2^+^ NSPCs among the total EGFP^+^ cells was unchanged (*p* = 0.290), whereas EGFP^+^Tbr2^+^ IP proportion was significantly increased relative to control (*p* = 0.028) (**Fig. 4f, g**). Subsequently, to examine *Pbdc1* overexpression (OE) effects, we delivered *pCAX-Pbdc1* vector at E14.5 and confirmed successful upregulation of *Pbdc1* mRNA at E15.5, 24 h post-IUE (**Supplementary Fig. 2b**). Similar to *Pbdc1*-KD, EGFP^+^Sox2^+^ NSPC population was unchanged (*p* = 0.973; **Fig. 4h**), whereas Tbr2^+^EGFP^+^ IP fraction was markedly decreased in *Pbdc1*-OE group (*p* = 6.07×10^-3^; **Fig. 4i**). Collectively, these results indicate that Pbdc1 restrains the size of Tbr2^+^ IP pool without grossly altering the Sox2^+^ NSPC population. The reduction of IPs in *Sey*/+ females along with elevated *Pbdc1* expression suggests Pax6- and/or sex-dependent transcriptional control of *Pbdc1* (see discussion).

### Female-biased H3K4me3 upregulation at the *Pbdc1* promoter in *Sey*/+ embryonic cortex

To investigate the mechanism underlying the *Sey*/+ female-specific upregulation of *Pbdc1*, we examined its transcriptional regulation in the developing cortex. Given the evidence that Pax6 cooperates with BAF (SWI/SNF) chromatin-remodeling complexes to repress downstream targets^39^, we hypothesized that Pax6-BAF axis regulates *Pbdc1* transcription. Chromatin immunoprecipitation (ChIP)-qPCR using E14.5 WT cortices demonstrated robust Pax6 occupancy at the *Pbdc1* promoter in both sexes, significantly above the negative control *Hoxd3* locus, with no sex differences (**Fig. 5a, left**). BAF components (Baf155 and Brg1) were likewise enriched at the *Pbdc1* promoter relative to *Hoxd3* in females (*p* = 1.19×10^-4^ and *p* = 0.035, respectively), with similar but non-significant trend in males (Baf155, *p* = 0.110; Brg1, *p* = 0.058; **Fig. 5a, middle and right**).

**Fig. 5:**
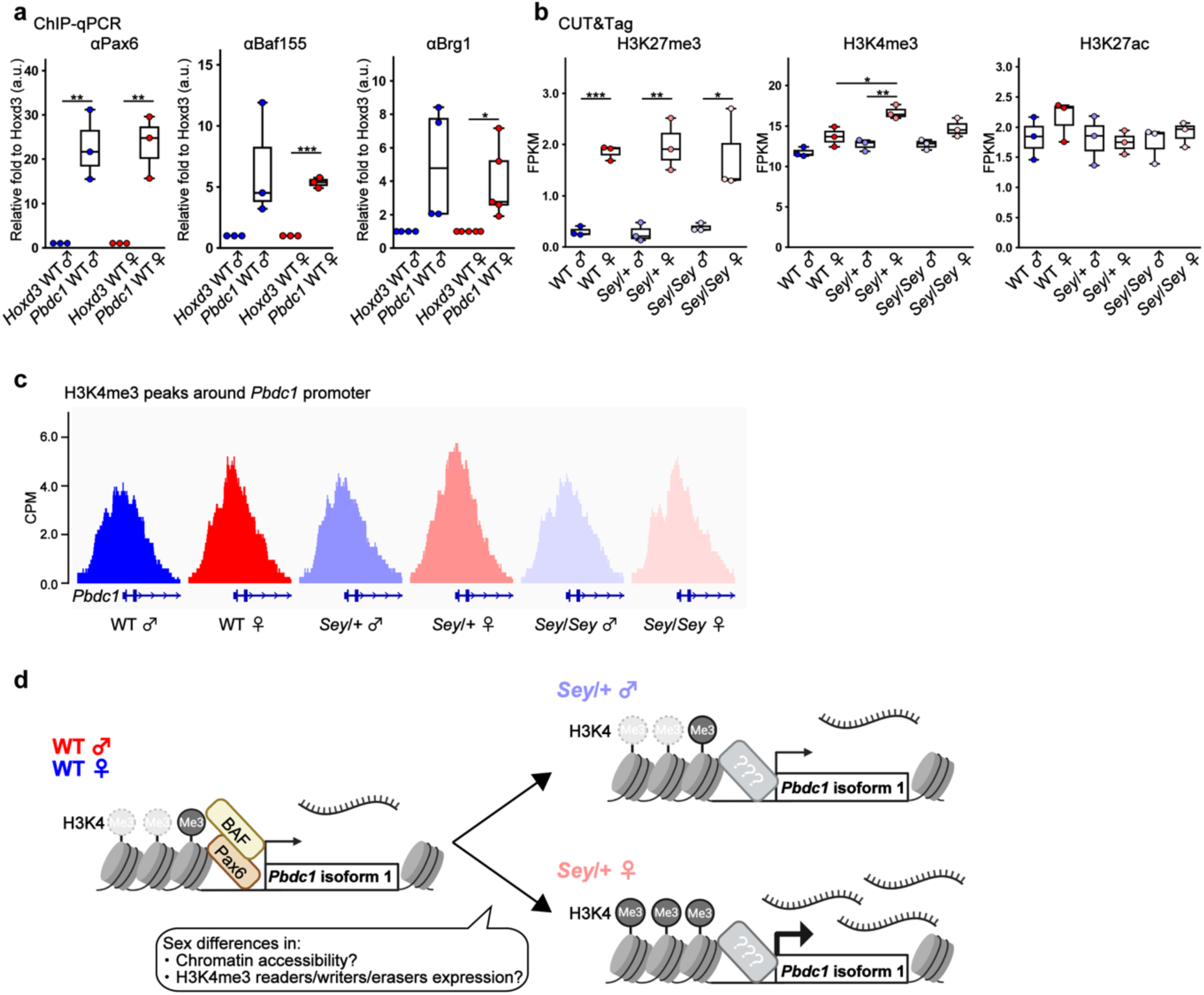
Sex differences in the deposition of H3K4me3 at the *Pbdc1* promoter. **a** ChIP-qPCR analysis of Pax6, Baf155 and Brg1 at the promoter of *Hoxd3* (control) and *Pbdc1* in WT male and female cortices. **b** Quantification of the CUT&Tag analysis on H3K27me3, H3K4me3 and H3K4ac at *Pbdc1* promoter in WT, *Sey*/+ and *Sey*/*Sey* male and female cortices. **c** IGV view showing H3K4me3 intensity at *Pbdc1* promoter locus in the embryonic cortices of WT, *Sey*/+ and *Sey*/*Sey* males and females. All 3 samples of each genotype/sex are merged. **d** Schematic illustration of hypothetical model proposing how *Pbdc1* transcription is regulated. \**p* < 0.05, \*\**p* < 0.01, \*\*\**p* < 0.001; determined by two-tailed Student’s *t*-test followed by Benjamini-Hochberg multiple testing correction. The line in the middle of box plots represent median, the lower and upper bounds of boxes indicate the first and third quartiles, and end of whiskers denote the minimum and maximum values. FPKM, fragments per kilobase of exon per million mapped reads; CPM, count per million.

We next profiled promoter-proximal histone modifications for H3K27me3, H3K4me3 and H3K4ac by cleavage under targets and tagmentation (CUT&Tag) in the cortices derived from WT, *Sey*/+ and *Sey*/*Sey* embryos of both sexes at E14.5. Quantitative analyses revealed higher H3K27me3 in females across all genotypes (WT males vs. WT females, *p* = 9.65×10^-5^; *Sey*/+ males vs. *Sey*/+ females, *p* = 5.55×10^-3^; *Sey*/*Sey* males vs. *Sey*/*Sey* females, *p* = 0.040; **Fig. 5b, left**), consistent with globally elevated H3K27me3 on female inactive X chromosome (Xi)^40^. Strikingly, H3K4me3, a mark associated with transcriptional activation, was selectively increased at the *Pbdc1* promoter in *Sey*/+ females relative to *Sey*/+ males (*p* = 3.66×10^-3^), whereas no sex difference was observed in WT or *Sey*/*Sey* samples (**Fig. 5b, middle; Fig. 5c**). By contrast, H3K27ac showed no detectable differences across genotypes or sexes (**Fig. 5b, right**). Together, these data indicate that the *Sey*/+ female-biased upregulation of *Pbdc1* is associated with enhanced promoter-proximal H3K4me3 and support a model in which Pax6, acting with BAF complexes, contributes to *Pbdc1* transcriptional regulation.

## Discussion

This study provides, to our knowledge, the first comprehensive analysis of sex differences in embryonic cortical gene expression and identifies a sex-biasing mechanism that cannot be explained solely by canonical X chromosome inactivation (XCI) escapees. In the *Pax6* haploinsufficient (*Sey*/+) telencephalon, sex differences in transcriptomes were markedly amplified relative to WT and *Sey*/*Sey*, and we identified *Pbdc1*, a previously uncharacterized X-linked gene, as a candidate regulator of sex-biased cortical development, with robust expression in NSPCs and IPs. Proximity-dependent PPI indicated that Pbdc1 associates with mRNA-splicing/processing factors, mainly implicated in cell-cycle control, while *in vivo*, perturbations demonstrated that Pbdc1 restrains the size of the IP pool without grossly altering the Sox2^+^ NSPC compartment. We further found Pax6 and BAF occupancy at the *Pbdc1* promoter in WT embryos of both sexes and a selective increase of promoter-proximal H3K4me3 in *Sey*/+ females, supporting a model in which Pax6-BAF normally constrains *Pbdc1* and Pax6 partial reduction unmasks a female-biased promoter activation state, as discussed below (**Fig. 5d**).

Sex-biased transcription is detectable in the WT mouse brains before perinatal sex-hormone exposure, with a strong contribution from sex-chromosome genes^41–43^. Consistent with our previous datasets^24,44^, WT E14.5 telencephalon again showed sex-DEGs that were largely enriched for X- and Y-linked loci. Notably, the *Sey*/+ background magnified these sex differences, extending earlier reports of sex-specific morphological alterations in the postnatal *Sey*/+ brains^23^ back to embryonic stages. In contrast, sex differences were attenuated in *Sey*/*Sey*, which might be due to a profound developmental arrest and altered cell-type composition resulting from the collapse of transcriptional programs under the complete loss of functional Pax6. Such a dosage-dependent effect is also evident in the other Pax6-dependent organogenesis, manifested as anophthalmia in *Sey*/*Sey* and a variety of ocular phenotypes in *Sey*/+, particularly on the C57BL/6 background.^45,46^.

Our isoform-resolved analyses clarify how *Pbdc1* fits within, and goes beyond, the classic “X-escapee” framework. Previous work reported female-biased *pan-Pbdc1* expression in the embryonic rat brain^47^ and escape from XCI in interspecific female cell lines^48,49^ and in adult murine tissues^50^, but these prior studies did not distinguish isoforms. In our data, isoform 2 accounted for the modest female bias in WT, consistent with XCI escape. In contrast, isoform 1, the isoform that increased most in *Sey*/+ females, showed comparable expression between WT males and WT females and thus cannot be explained by XCI escape alone. This underscores the importance of isoform-level resolution when interrogating sex-biased programs in a setting where alternative splicing is pervasive^51^.

Functional clues came from Pbdc1’s interactome in a mammalian neuronal context. Contrary to hypothesis stemming from non-mammalian homologs linked to transcriptional machinery^52–54^, AirID based high-resolution proximity-dependent PPI analysis in N2 cells revealed preferential interaction with RNA-splicing and processing factors, over half of which have established roles in proliferation and oncogenesis, such as HNRNPK^33^, SNRPA1^37^. and SART1^35^. Together with GO shifts in *Sey*/+ (female-biased upregulation of RNA processing genes; male-biased enrichment of developmental programs), these findings support a model in which Pbdc1 modulates RNA-splicing networks that gate proliferative capacity and fate transitions during corticogenesis.

*In vivo* manipulations were consistent with this model. *Pbdc1*-KD increased Tbr2^+^ IPs, whereas *Pbdc1*-OE decreased them, without changing Sox2^+^ NSPC population. Because murine cortical neurons are produced either directly from NSPCs or indirectly via IPs during murine corticogenesis and ∼78% of them arise via the IP population^55^, the IP reduction upon *Pbdc1*- OE suggests that Pbdc1 either (i) restricts NSPC commitment to the IP lineage, and/or (ii) accelerates IP-to-neuron differentiation, shortening the transit- amplifying window. Distinguishing these possibilities will require lineage tracing, birth-dating, and kinetic analyses of IP proliferation and neurogenic divisions. The reduced IP population in *Sey*/+ females, with elevated *Pbdc1* expression, parsimoniously link *Pbdc1* upregulation to sex-biased cytoarchitectural outcomes.

Mechanistically, we propose a two-factor model for Pbdc1 regulation (**Fig. 5d**). First, in WT, Pax6 with BAF co-occupies the *Pbdc1* promoter and restrains its transcription, consistent with prior evidence of Pax6-BAF-mediated repression and the elevated *Pbdc1* (shown as *2610029G23Rik*) reported in *Sey*/*Sey* cortices (sex unspecified)^56^. Second, partial Pax6 reduction in *Sey*/+ reveals a female-biased promotor activation state, reflected by increased H3K4me3 specifically in *Sey*/+ females, while H3K27ac remains unchanged. We note that H3K27me3 was globally higher in females across genotypes, consistent with Xi-linked elevations only in females^40^, but only H3K4me3 at *Pbdc1* promoter tracked the *Sey*/+ female-specific transcriptional increase. A plausible contributor is sex-biased dosage and activity of H3K4 methyl readers/writers/erases: female-biased *Kdm5c* (X-linked) and male-biased *Kdm5d* (Y-linked) demethylases^24,41–43^ and/or sex-biased chromatin accessibility among male X chromosome and female active X chromosome^57^. Those factors could differentially tune promoter H3K4me3 when Pax6-mediated repression is weakened.

Limitations of our study include: (i) modest RNA-seq sample size (n = 3 per group), which can inflate false positives for small effect-size of sex differences^58^, hence our use of orthogonal validations (RT-qPCR and RNAscope); (ii) lack of a validated anti-Pbdc1 antibody precluded protein-level quantitation; (iii) AirID was performed in N2a cells rather than embryonic cortex, warranting *in situ* confirmation of its interactome (e.g., IP-wester blotting or proximity labeling *in vivo*); and (iv) we did not directly demonstrate Pbdc1-dependent splicing changes in NSPCs/IPs. Finally, the connection between Pbdc1 dysregulation and ASD-relevant phenotypes remains inferential.

Despite these caveats, our work establishes a conceptual advance: sex-biased transcriptional architecture in the embryonic cortex can be potentiated by a dosage-sensitive transcription factor (Pax6) that gates a female-preferential promoter activation state (H3K4me3) at a previously uncharacterized X-linked gene (*Pbdc1* isoform 1), a mechanisms distinct from canonical XCI escape. This framework motivates further genetic tests (e.g., isoform-specific knock-in/overexpression of *Pbdc1* via CRISPR-mediated technology) paired with single-cell transcriptomics and splicing-aware analyses to delineate Pbdc1-centered regulatory circuits and access their impact on neurodevelopmental trajectories and behavior.

## Method

### Experimental animals

All animal experiments were conducted according to the National Institutes of Health Guide for the Care and Use of Laboratory Animals and were approved by the Ethics Committee for Animal Experiment of Tohoku University Graduate School of Medicine (approved no. 2020MdA-109-20) and by the Animal Care and Use Committee of The University of Tokyo Institute for Quantitative Biosciences (approved no. A2022IQB001). Male and female WT (C57BL/6 J) mice were purchased from CLEA Japan and used to maintain the colony of *Pax6* mutant (*Sey*) mice^59^ at the animal facility of Tohoku University Graduate School of Medicine. Mice were housed under a 12-h light/dark cycle (lights on at 8:00 and off at 20:00) with *ad libitum* access to food and water. Embryonic day 0.5 (E0.5) was defined as noon on the day when a vaginal plug was detected.

### DNA extraction and genotyping for *Ube1* and *Sey*

Genomic deoxyribonucleic acid (DNA) was extracted from the tail tissue of the embryos. A mixture of 10 μl of 5x Colorless GoTaq® Flexi Buffer (Promega), 5 μl of 10% NP-40 (Merck Millipore), 2 μl of Proteinase K (Sigma; 20 mg/ml) and 33 μl of Milli-Q water was added to the tubes with mouse tissue and incubated overnight at 57 °C on a platform shaker. The samples were then heated at 90-95 °C for 15 min in water bath and centrifuged at 15,000 rpm for 15 min at 4 °C. The supernatant (crude DNA extract) was stored at 4 °C. Polymerase chain reaction (PCR) was performed using 1 μl of the DNA extract and the GoTaq® Flexi DNA Polymerase (Promega). Sex determination (*Ube1*) was performed using forward primer for *Ube1*-F 5′- TGGTCTGGACCCAAACGCTGTC-3′ and reverse primer for *Ube1*-R 5′- GGCAGCAGCCATCACATAATCC-3′^60^. The *Pax6 Sey* allele was genotyped using two forward primers (SP1: 5′-GAGAACACCAACTCCATCAGTTCTAAGT-3′; SP2: 5′- AGCAACAGGAAGGAGGGGGAACGAACACCAACTCCATCAGTTCTTACG-3′) and one reverse primer (MC130: 5’-CTTTCTCCAGAGCCTCAATCTG-3’)^61^. PCR was performed using the Vapo Protect Thermal Cycler (Eppendorf) as follows: For *Ube1*: 95 °C for 2 min, followed by 40 cycles of 95 °C for 15 sec, 60 °C for 20 sec, and 72 °C for 1 min, with a final extension at 72 °C for 2 min. For *Sey,* an initial denaturing at 95°C for 2 min, annealing at 55°C for 1 min, and extension at 72°C for 1 min, followed by 30 cycles of 95°C for 40 sec, 55°C for 40 sec, and 72°C for 40 sec, with a final extension at 72°C for 10 min. The amplified PCR products were visualized by electrophoresis on 3% agarose gels using the Gel DocTM EZ Imager (Bio-Rad).

### RNA-seq

As *Sey*/*Sey* mice are perinatally lethal, *Sey*/+ male and female mice were intercrossed to obtain six genotypes (WT male, WT female, *Sey*/+ male, *Sey*/+ female, *Sey*/*Sey* male, *Sey*/*Sey* female). At E14.5, the whole telencephalons were precisely dissected using fine forceps and micro- scissors after carefully removing the overlying skin and skull, following previous methods^24,62^. The dissected tissue was immediately homogenized in Buffer RLT Plus (RNeasy Plus Mini Kit, QIAGEN) supplemented with β-mercaptoethanol and stored at −80 °C. Tail samples were simultaneously collected from the embryos for sex determination and *Sey* genotyping using PCR (refer to “DNA extraction and genotyping for *Ube1* and *Sey*” section). For RNA extraction, three biological replicates were prepared for each genotype. Total RNA was extracted from the telencephalons using the RNeasy Plus Mini Kit (QIAGEN). Sequencing of extracted RNA for a total of 18 samples was commissioned by Takara Bio Inc. cDNA libraries were generated using TruSeq Stranded mRNA Sample Prep Kit (Illumina) on XT-Auto System (Agilent), and subsequently sequenced on NovaSeq 6000 platform (Illumina) with 150 bp paired-end modules. All submitted RNA samples for sequencing displayed high integrity with RIN value ≥ 9.8.

### RNA-seq data analysis

Raw RNA-seq data were processed utilizing the Rhelixa RNA-seq pipeline (https://sc.ddbj.nig.ac.jp/en/advanced_guides/Rhelixa_RNAseq/Rhelixa_RNAseq) implemented on the NIG supercomputer system. The workflow comprised the following data processing steps:

1. Assessment of the quality using FastQC v0.11.7 (--nogroup) (https://www.bioinformatics.babraham.ac.uk/projects/fastqc/)^63^.
2. Removal of adaptors and quality trimming using Trimmomatic v0.3826^64^ (ILLUMINACLIP:paired_end.fa:2:30:10, LEADING:20, TRAILING:20, SLIDINGWINDOW:4:15, MINLEN:36) with both Illumina standard and custom adapter sequences.
3. Assessment of strand information with RSeQC v3.0.127^65^ (infer_experiment.py) using the mm10 RefSeq Gene annotation.
4. Alignment of reads to the mm10 reference genome with HISAT2 v2.1.028^66^ (--rna-strandness RF, --dta).
5. Conversion to BAM file and sorting using SAMtools v1.929^67^ (view, sort, index).
6. Quantification of reads with featureCounts v1.6.330^68^ (-p, -s 2, -T 4, -F GTF, -t exon, -g gene_id) against the Ensembl Mus_musculus.GRCm38.87 annotation.

The resulting raw RNA-seq read count table was imported into R (v4.3.3) for differential expression analysis using edgeR (v4.0.16)^69^. For each pairwise comparison (WT male vs. female; *Sey*/+ male vs. female; *Sey*/*Sey* males vs. females), appropriate columns from the read count table were selected and an intercept design matrix was created using the model. The count matrix for gene-level quantification is provided with the result of DE analysis (**Supplementary Table 2**).

In parallel to the steps above, isoform-level quantification of the RNA-seq was carried out utilizing *Salmon* (v1.10.3)^29^. The resulting isoform-level count table was also imported into R for differentially expressed transcripts analysis using edgeR^69^. The count matrix for isoform-level quantification is provided with the result of DEG analysis (**Supplementary Table 3**). Hierarchical clustering and heatmap was generated using Python (v3.9.18). Additionally, GO analyses for Biological Process were conducted with ShinyGO^70^.

### RT-qPCR

Telencephalon samples were collected from embryo at E14.5. Total RNA was extracted with the RNeasy Mini Kit (QIAGEN) according to the manufacturer’s instructions. Subsequently, reverse transcription (RT) was performed with SuperScript™III First-Strand Synthesis System (Invitrogen). RT-qPCR was performed as previously described^71^, in technical duplicate, using TaqPro Universal SYBR qPCR Master Mix (Vazyme) and CFX96 Real-Time PCR Detection System (Bio-Rad). The relative expression levels of each target were calculated employing 2−ΔΔCt method, with *Sdha* serving as the internal control^72^. Primers were designed with Primer3Plus^73^, and detailed primer sequences are provided in **Supplementary Table 4**

### *in situ* hybridization

The embryos were fixed by transcardial perfusion with 4% paraformaldehyde (PFA) dissolved in phosphate-buffered saline (PBS). The brains were post-fixed overnight in 4% PFA in PBS at 4°C and cryoprotected sequentially in 10% and 20% sucrose until brains sank. Brains were then embedded in Tissue-Tek OCT compound (Sakura). *in situ* hybridization on frozen coronal sections was performed as previously described^74^ with frozen coronal sections (14 µm) sliced with a cryostat (Leica, CM3050S). For probe preparation, RNA was extracted from E14.5 WT telencephalons using RNeasy Plus Mini Kit (QIAGEN), and cDNA was synthesized with the SuperScript™III First-Strand Synthesis System (Invitrogen). The cDNA fragments of *Pbdc1* (NM_026312, nucleotides 1-467, this sequence is completely identical with nucleotides 1-467 of NM_001281871, isoform 2) were cloned into pBluescript II SK (–) (Stratagene). The DIG-labeled RNA probes were generated from these plasmids using the DIG RNA Labeling kit (Roche). Hybridization signals were visualized using a fluorescence microscope (BZ-X710, KEYENCE).

### RNA-scope

The embryos were fixed by transcardial perfusion with 4% PFA in PBS. The brains were post-fixed overnight in 4% PFA in PBS at 4°C and cryoprotected sequentially in 10% and 20% sucrose until brains sank. Brains were then embedded in Tissu-Tek OCT compound (Sakura). Coronal sections (14 µm) were cut on a cryostat (Leica, CM3050S) and completely dried in the chamber of cryostat for 1 h at −22 °C. The Probe for *Pbdc1* isoform 1 was designed by the manufacturer and detection of the mRNA was performed using RNAscope® Fluorescent Multiplex Reagent Kit (ACD, 320850) according to the manufacturer’s protocol. All images were acquired using a confocal laser scanning microscope Zeiss LSM800 (Carl Zeiss).

### *In utero* electroporation

*In utero* electroporation (IUE) was performed as previously described^75^. Pregnant WT mice at E14.5 were deeply anesthetized with isoflurane. A solution containing siRNA (final concentration of 2 µg/µL; *Pbdc1* isoform 1 RNAi: UAAUCCUGAUACCAGGUACAAGCUA; Scramble control RNAi: UAAGAUCCGUUACAACGGAUACCUA), *pCAX-Pbdc1* (NM_026312.5, isoform 1; final concentration of 2 µg/µL) were injected into the lateral ventricle of the embryonic brains by using a glass capillary with the electric microinjector system (IM-300, Narishige) together with *pCAG-EGFP* (final concentration of 0.5 μg/μl) and Fast green dye. Electroporation was conducted by applying five electric pulses at 40 V, each lasting 50 ms, at 950 ms intervals, through the uterus using forceps-type electrodes (LF650P5, BEX) connected to an electric gene transfer device (CUY-21, BEX).

### Immunohistochemistry

Immunohistochemistry was performed as previously described^74^. The embryos were fixed by transcardial perfusion with 1% PFA in PBS. The brains were post-fixed overnight in 1% PFA in PBS at 4°C and cryoprotected sequentially in 10% and 20% sucrose until brains sank. Brains were then embedded in Tissu-Tek OCT compound (Sakura). Coronal sections (14 µm) were cut on a cryostat (Leica, CM3050S). The frozen sections were incubated overnight at 4℃ with primary antibodies diluted in blocking solution (3% BSA in TBS with 0.1% Triton X-100 (TBST). On the following day, the sections were rinsed three times in TBST and subsequently incubated for 1 h at room temperature (RT) with fluorophore-conjugated secondary antibodies and 4′,6-Diamidino-2-phenylindole dihydrochloride (DAPI, 1:1000, Sigma). After three further washes in TBST, the sections were mounted with VECTASHIELD® antifade mounting medium and covered with coverslips.

For double staining of mRNA (ISH) and protein (immunohistochemistry), sections were washed in TBST for 10 min, followed by PBS for 10 min after color reaction step of ISH. Sections were subsequently fixed in 4% PFA in PBS for 10 min at RT, and then washed in PBS for 5 min and in TBST for 5 min twice. The subsequent procedures were performed as described in this “Immunohistochemistry” section, starting from the blocking step. Antibodies used are listed in the **Supplementary table 5.** All images were acquired using a confocal laser scanning microscope Zeiss LSM800 (Carl Zeiss).

### Cell culture and generation of stable cell lines using lentivirus

HEK293T (CRL-3216, American Type Culture Collection [ATCC], RRID:CVCL_0063) cells were cultured in low-glucose DMEM (DMEM, 041-29775; FUJIFILM Wako) supplemented with 10% foetal bovine serum (535-94155, FUJIFILM Wako), 100 µg/mL penicillin, and 100 µg/mL streptomycin (15140122, Thermo Fisher Scientific, MA, USA) at 37 °C under 5% CO2. HEK293T cells were transiently transfected with TransIT-LT1 transfection reagent (MIR2304, Mirus Bio). N2a cells were cultured in RPMI 160 GlutaMAX medium (72400047, Gibco) supplemented with 10% foetal bovine serum (535-94155, FUJIFILM Wako), 1 mM Sodium Pyruvate (11360070, Gibco) and 25 mM HEPES at 37 °C under 5% CO2.

Each lentivirus was produced in HEK293T cells by transfection with the pCSII-CMV-ORF- IRES2-Bsd expression vector together with pCMV-VSV-G-RSV-Rev and pCAG-HIVgp as described in previous study^76^. N2a cells supplemented with 10 µg/mL polybrene (12996-81, Nacalai Tesque) were infected with the appropriate lentivirus. After 24 h of infection, the culture medium was replaced, and 10 µg/mL blasticidin S (ant-bl-10p, InvivoGen) selection was started 24 h thereafter.

### Mass spectrometry analysis of biotinylated peptides

Proximity-dependent PPI analysis using AirID was performed essentially as described previously^32,77^, with modifications. N2a cell stably expressing AGIA-AirID-Pbdc1 isoform 1 or AGIA-AirID-IκBα were cultured in 10-cm dishes and incubated with 10 μM biotin for 6 h. After washing three times with 10 mL HEPES saline buffer (20 mM HEPES-NaOH, pH 7.5, 137 mM NaCl), the cells were lysed in 300 µL Gdm-TCEP buffer (6 M guanidine-HCl, 100 mM HEPES-NaOH [pH 7.5], 10 mM TCEP, 40 mM chloroacetamide). Cell lysates were pooled in one tube and divided into three tubes of 300 µL each, followed by digestion with trypsin. The resulting peptides were incubated with Tamavidin 2-Rev magnetic beads (FUJIFILM Wako). Biotinylated peptides were eluted with 1 mM biotin prior to mass spectrometry analysis.

LC-MS/MS analysis of the peptides was performed using an EASY-nLC 1200 UHPLC system coupled to an Orbitrap Fusion mass spectrometer (Thermo Fisher Scientific) with a nanoelectrospray ion source. Separation of the peptides was performed on a 75-µm inner diameter × 150 mm C18 reversed-phase column (Nikkyo Technos) with a linear gradient of 4– 32% acetonitrile over 60 min, followed by a ramp to 80% acetonitrile for 10 min and a final hold at 80% acetonitrile for 10 min. The mass spectrometer was operated in data-dependent acquisition mode with a maximum duty cycle for 3 sec. Full MS scans (MS1) were acquired at a resolution of 120,000 (at m/z 200), AGC target 4e5, mass range 375–1500 m/z. HCD MS/MS spectra (MS2) were acquired in a linear ion trap with AGC target 1e4, isolation window 1.6 m/z, normalized collision energy 30, maximum injection time 200 ms, and dynamic exclusion of 10 sec. Raw MS data were analyzed using Proteome Discoverer version 2.5 (Thermo Fisher Scientific) against the SwissProt mouse database. Search parameters included: enzyme, trypsin (up to two missed cleavages); precursor mass tolerance, 10 ppm; fragment mass tolerance, 0.6 Da; fixed modification, carbamidomethylation of cysteine; variable modifications, protein N-terminal acetylation, methionine oxidation, and lysine biotinylation. Peptide identification was filtered at a false discovery rate (FDR) of 1% using the Percolator node. Quantification was performed by label-free precursor ion intensities, and normalization was applied to equalize the total peptide abundance across samples (**Supplementary Table 6**). Additionally, GO analyses (Biological Process) of interacting proteins were conducted with ShinyGO^70^. The proteins interacting with Pbdc1 and implicated in RNA-splicing terms are listed in **Supplementary Table 1** with references indicating the function of interactors in cell-cycle regulation or cancer cell proliferation^33–35,78–103^.

### ChIP-qPCR

ChIP for Pax6, Baf155 and Brg1 was performed as previously described^104^with minor modifications. Embryonic dorsal cortices were dissociated with by Papain Dissociation System (Worthington) as described previously^105^, followed by fixation with 1% formaldehyde in PBS and immediately frozen in liquid nitrogen, and stored at –80 °C. After thawing, the cells (approximately 2 million cells) were resuspended in RIPA buffer (10 mM Tris HCl at pH 8.0, 1 mM EDTA, 140 mM NaCl, 1% Triton X 100, 0.1% SDS, and 0.1% sodium deoxycholate) and sonicated using a Picoruptor (15 cycles of 30 s ON and 30 s OFF) (Diagenode). Resulting cell lysates were diluted with RIPA buffer for immunoprecipitation (50 mM Tris-HCl at pH 8.0, 150 mM NaCl, 2 mM EDTA, 1% Nonidet P 40, 0.1% SDS, and 0.5% sodium deoxycholate) and incubated for 1 h at 4 °C with Protein A/G Magnetic Beads (Pierce) to avoid non-specific reactivity. Subsequently, they were incubated overnight at 4 °C with Protein A/G Magnetic Beads that had previously been incubated for overnight at 4 °C with antibodies. The beads were isolated and rinsed first three times with wash buffer (2 mM EDTA, 150 mM NaCl, 0.1% SDS, 1% Triton X100, and 20 mM Tris HCl at pH 8.0) and then once with wash buffer containing 500 mM NaCl. Immune complexes were eluted from the beads for 15 min at 65 °C with a solution containing 10 mM Tris HCl at pH 8.0, 5 mM EDTA, 300 mM NaCl, and 0.5% SDS, and they were subjected to digestion with proteinase K (Nacalai) for >6 h at 37 °C, removal of cross links by incubation at 65 °C for >6 h, and extraction of the DNA leftover with phenol– chloroform–isoamyl alcohol and ethanol. Purified DNA was washed with 70% ethanol, suspended in water, and subjected to real-time qPCR analysis in a LightCycler 480 (Roche) with Thunderbird SYBR qPCR Mix (Toyobo). Antibodies and primer sequences are provided in **Supplementary Table 4** and **5.**

### CUT&Tag sequence

CUT&Tag was conducted as previously described^106^, with minor modifications. Embryonic dorsal cortices were isolated from embryos and dissociated by Papain Dissociation System (Worthington). Dissociated cells were dispersed in CELLBANKER 1 (ZENOAQ), and stored at –80 °C until use. Concanavalin A beads (Bangs Laboratories) were washed with binding buffer (10 mM KCl, 1 mM CaCl2, 1 mM MgCl2, and 20 mM HEPES at pH 7.5), mixed with thawed cells and rotated for 10 to 20 min at RT. After washing with Wash buffer (150 mM NaCl, 0.5 mM spermidine, a protease inhibitor (Roche) and 20 mM HEPES at pH 7.5) once, the beads were suspended in Antibody buffer (150 mM NaCl, 0.5 mM spermidine, a protease inhibitor, 0.05% digitonin, 2 mM EDTA, 0.1% BSA, and 20 mM HEPES at pH 7.5) with primary antibodies for overnight at 4 °C. After washed with Dig-wash buffer (150 mM NaCl, 0.5 mM spermidine, a protease inhibitor, 0.05% digitonin, and 20 mM HEPES at pH 7.5) once, the beads were suspended in Dig-wash buffer with secondary antibodies for 30 min at RT. The beads were washed with Dig-wash buffer for three times and subjected to binding reaction of pAG-Tn5 adaptor complex in Dig-300 buffer (300 mM NaCl, 0.5 mM spermidine, protease inhibitor, 0.01% digitonin, and 20 mM HEPES at pH 7.5) for 1 h at RT. Note that pAG-Tn5 was constructed with the usage of both pAG-MNase (Addgene #123461) and pA-Tn5 (Addgene #124601) plasmids and was used instead of pA-Tn5 in the original protocol. After washed with Dig-300 buffer for three times, the beads were suspended with Tagmentation buffer (300 mM NaCl, 0.5 mM spermidine, a protease inhibitor, 0.01% digitonin, 1 mM MgCl2, and 20 mM HEPES at pH 7.5) for tagmentation reaction for 1 h at 37 °C. The tagmented DNA was eluted by adding 0.5 M EDTA (final 17 mM), 10% SDS (final 0.1%), and 10 mg/ml proteinase K (final 0.17 mg/ml) for overnight at 37 °C, purified using phenol–chloroform–isoamyl alcohol and ethanol, washed with 70% ethanol and suspended in water. The DNA was amplified by PCR reaction using Q5 Hot Start High-Fidelity 2x Master Mix (NEB) as follows: 5 min at 72 °C and 30 s at 98 °C followed by 12 cycles of 10 s at 98 °C and 20 s at 63 °C, with a final extension at 72 °C for 1 min. Finally, the amplified DNA was purified using SPRI beads (Beckman Coulter). Antibodies are listed in **Supplementary Table 5**.

The libraries for CUT&Tag were sequenced as 36-bp paired-end reads on the Illumina Nextseq2000 platforms. Approximately 20 million sequences were obtained from each sample. The reads were subsequently mapped to a mouse genome (mm10) using bowtie2^107^. Among the uniquely mapped reads, the reads in the blacklist regions determined by the Encyclopedia of DNA Elements project^108,109^, and duplicated reads were removed using samtools rmdup function^67^. The signals in the promoter regions were quantified using BAMscale^110^. Aligned reads were monitored with Integrative Genomics Viewer (IGV)^111^.

### Statistical analysis

Statistical details for each experiment are provided in corresponding figure legends. Data are presented as boxplots, where the line in the middle of boxes represent median; the lower and upper bounds of boxes indicate the first and third quartiles, and end of whiskers denote the minimum and maximum values. Student’s *t*-test, followed by Benjamini-Hochberg multiple testing was carried out by Python (v 3.9.18).

## Data availability

All public data and materials used in the analysis are cited and referenced in the manuscript. The raw read sequences of bulk RNA-seq are available in DNA Databank of Japan (DDBJ) under the accession number of PRJDB37737. Count table of RNA-seq is available as **Supplementary Table 2** (gene-level quantification) and **3** (isoform-level quantification). Proximity-dependent PPI data is available as **Supplementary Table 6**. The raw read sequences and processed data of CUT&Tag analyses are available in DNA Databank of Japan (DDBJ) under the accession numbers of PRJDB37690.

## Code availability

All publicly available software tools used in the analyses are cited and referenced in the manuscript.

## Acknowledgements

We thank Dr. Satoshi Miyashita (National Center of Biochemistry and Cellular Biology), Dr. Kazumichi Fujiwara (National Institute of Genetics) and Dr. Yoshio Wakamatsu (The University of Melbourne) for technical support in bioinformatical analyses and for valuable comments. N2a cell was kindly provided by Dr. Kenji Iemura and Prof. Kozo Tanaka (Tohoku University). We are grateful to Ms. Sayaka Makino and Ms. Asuka Takehara for animal care and technical assistance, and to all members of the Osumi Laboratory for helpful discussions. This study was supported by JSPS KAKENHI (#23KJ0193 to S.M.; #21K15097 and #24H01419 to S.O.; #24K02203 to T.K.), JST SPRING (#JPMJSP2114 to S.M.), AMED (#JP24wm0625311 to T.K.; #JP21wm0425003 to N.O.). We also acknowledge “Support System for Young Researchers to Use Research Equipment, Instruments and Devices” at Tohoku University. Computations were partially performed on the NIG supercomputer at ROIS National Institute of Genetics. Additional support was provided by Molecular Profiling Committee, Grant-in-Aid for Transformative Research Areas “Advanced Animal Model Support (AdAMS)” from MEXT/JSPS (#JP22H04922) and Medical Research Center Initiative for High Depth Omics (to H.K.).

## Author contributions

S.M., S.O., T.K. and N.O. designed the study; S.M. and S.O. collected samples for RNA-seq; S.M. analyzed RNA-seq data and performed RT-qPCR, *in situ* hybridization, immunohistochemistry and following image analyses; S.M. and T.K. performed RNA-scope; S.M., S.N. and T.K. performed *in utero* electroporation; K.Y. and T.S. performed proximity-dependent PPI assay; H.K. performed mass spectrometry analysis; S.M. and S.O. performed tissue dissociation for ChIP-qPCR and CUT&Tag; M.S. and Y.K. performed ChIP-qPCR and CUT&Tag and the data analyses; S.M. drafted manuscript; S.O. T.K. and N.O. edited the manuscript; S.M., S.O., T.K. and N.O. acquired funding; All authors discussed the results and commented on the manuscript.

## References

1. Turano, A., Osborne, B. F. & Schwarz, J. M. Sexual Differentiation and Sex Differences in Neural Development. in Current Topics in Behavioral Neurosciences vol. 43 (2019).

2. Fujiyama, T. et al. Forebrain Ptf1a Is Required for Sexual Differentiation of the Brain. Cell Rep 24, (2018).

3. Pinares-Garcia, P., Stratikopoulos, M., Zagato, A., Loke, H. & Lee, J. Sex: A significant risk factor for neurodevelopmental and neurodegenerative disorders. Brain Sci 8, (2018).

4. Scharf, J. M., et al. Population prevalence of Tourette syndrome: A systematic review and meta-analysis. Movement Disorders 30, (2015).

5. Babinski, D. E. Sex Differences in ADHD: Review and Priorities for Future Research. Curr Psychiatry Rep 26, (2024).

6. 6. Maenner, M. J., et al. Prevalence and Characteristics of Autism Spectrum Disorder Among Children Aged 8 Years — Autism and Developmental Disabilities Monitoring Network, 11 Sites, United States, 2020. MMWR Surveillance Summaries 72, (2023).

7. Mandy, W. et al. Sex differences in autism spectrum disorder: Evidence from a large sample of children and adolescents. J Autism Dev Disord 42, (2012).

8. Werling, D. M. & Geschwind, D. H. Sex differences in autism spectrum disorders. Curr Opin Neurol 26, (2013).

9. Abrahams, B. S. et al. SFARI Gene 2.0: A community-driven knowledgebase for the autism spectrum disorders (ASDs). Mol Autism 4, (2013).

10. Walther, C. & Gruss, P. Pax-6, a murine paired box gene, is expressed in the developing CNS. Development 113, (1991).

11. Haubst, N. et al. Molecular dissection of Pax6 function: The specific roles of the paired domain and homeodomain in brain development. Development 131, (2004).

12. Manuel, M. N., Mi, D., Masonand, J. O. & Price, D. J. Regulation of cerebral cortical neurogenesis by the Pax6 transcription factor. Front Cell Neurosci 9, (2015).

13. Osumi, N., Shinohara, H., Numayama-Tsuruta, K. & Maekawa, M. Concise Review: Pax6 Transcription Factor Contributes to both Embryonic and Adult Neurogenesis as a Multifunctional Regulator. Stem Cells 26, (2008).

14. Kikkawa, T. et al. The role of Pax6 in brain development and its impact on pathogenesis of autism spectrum disorder. Brain Res 1705, (2019).

15. Ochi, S., Manabe, S., Kikkawa, T. & Osumi, N. Thirty Years’ History since the Discovery of Pax6: From Central Nervous System Development to Neurodevelopmental Disorders. Int J Mol Sci 23, (2022).

16. Gessler, M., Simola, K. O. J. & Bruns, G. A. P. Cloning of breakpoints of a chromosome translocation identifies the AN2 locus. Science (1979) 244, (1989).

17. Xu, S. et al. Characterization of 11p14-p12 deletion in WAGR syndrome by array CGH for identifying genes contributing to mental retardation and autism. Cytogenet Genome Res 122, (2008).

18. Davis, L. K. et al. Pax6 3′ deletion results in aniridia, autism and mental retardation. Hum Genet 123, (2008).

19. Maekawa, M. et al. A novel missense mutation (Leu46Val) of PAX6 found in an autistic patient. Neurosci Lett 462, (2009).

20. Umeda, T. et al. Evaluation of pax6 mutant rat as a model for autism. PLoS One 5, (2010).

21. Yoshizaki, K. et al. Paternal aging affects behavior in Pax6 mutant mice: A gene/environment interaction in understanding neurodevelopmental disorders. PLoS One 11, (2016).

22. Hiraoka, K. et al. Regional volume decreases in the brain of Pax6 heterozygous mutant rats: MRI deformation-based morphometry. PLoS One 11, (2016).

23. Joko, N., Kikkawa, T., Inoue, T. & Osumi, N. Sex-Difference in Olfactory Interhemispheric Malformation Caused by Pax6 Haploinsufficiency. Tohoku J Exp Med 266, 361–369 (2025).

24. Ochi, S. et al. A Transcriptomic Dataset of Embryonic Murine Telencephalon. Sci Data 11, 586 (2024).

25. Yuen, R. K. C. et al. Whole genome sequencing resource identifies 18 new candidate genes for autism spectrum disorder. Nat Neurosci 20, (2017).

26. Casellas-Vidal, D. et al. ZDHHC15 as a candidate gene for autism spectrum disorder. Am J Med Genet A 191, (2023).

27. Li, J. et al. Integrated systems analysis reveals a molecular network underlying autism spectrum disorders. Mol Syst Biol 10, (2014).

28. Luo, Y. et al. A multidimensional precision medicine approach identifies an autism subtype characterized by dyslipidemia. Nat Med 26, (2020).

29. Patro, R., Duggal, G., Love, M. I., Irizarry, R. A. & Kingsford, C. Salmon provides fast and bias-aware quantification of transcript expression. Nat Methods 14, (2017).

30. Loo, L. et al. Single-cell transcriptomic analysis of mouse neocortical development. Nat Commun 10, (2019).

31. Telley, L. et al. Temporal patterning of apical progenitors and their daughter neurons in the developing neocortex. Science (1979) 364, (2019).

32. Kido, K. et al. Airid, a novel proximity biotinylation enzyme, for analysis of protein– protein interactions. Elife 9, (2020).

33. Li, D., Guo, J. & Jia, R. Epigenetic Control of Cancer Cell Proliferation and Cell Cycle Progression by HNRNPK via Promoting Exon 4 Inclusion of Histone Code Reader SPIN1. J Mol Biol 435, (2023).

34. Kim, Y. D. et al. The unique spliceosome signature of human pluripotent stem cells is mediated by SNRPA1, SNRPD1, and PNN. Stem Cell Res 22, (2017).

35. Ozaki, K. et al. Involvement of the splicing factor SART1 in the BRCA1-dependent homologous recombination repair of DNA double-strand breaks. Sci Rep 14, 18455 (2024).

36. Stoykova, A., Fritsch, R., Walther, C. & Gruss, P. Forebrain patterning defects in Small eye mutant mice. Development 122, (1996).

37. Heins, N. et al. Glial cells generate neurons: The role of the transcription factor Pax6. Nat Neurosci 5, (2002).

38. Quinn, J. C. et al. Pax6 controls cerebral cortical cell number by regulating exit from the cell cycle and specifies cortical cell identity by a cell autonomous mechanism. Dev Biol 302, (2007).

39. Tuoc, T. C. et al. Chromatin Regulation by BAF170 Controls Cerebral Cortical Size and Thickness. Dev Cell 25, (2013).

40. Fang, H., Disteche, C. M. & Berletch, J. B. X Inactivation and Escape: Epigenetic and Structural Features. Front Cell Dev Biol 7, (2019).

41. Kuschel, B. et al. Sex differences in sex chromosome gene expression in mouse brain. Hum Mol Genet 11, (2002).

42. Dewing, P., Shi, T., Horvath, S. & Vilain, E. Sexually dimorphic gene expression in mouse brain precedes gonadal differentiation. Molecular Brain Research 118, (2003).

43. Szakats, S., McAtamney, A., Cross, H. & Wilson, M. J. Sex-biased gene and microRNA expression in the developing mouse brain is associated with neurodevelopmental functions and neurological phenotypes. Biol Sex Differ 14, (2023).

44. Ebrahimiazar, S. et al. A Transcriptomic Dataset of Embryonic Murine Telencephalon of Fmr1-Deficient Mice. Sci Data 12, 927 (2025).

45. Kanakubo, S. et al. Abnormal migration and distribution of neural crest cells in Pax6 heterozygous mutant eye, a model for human eye diseases. Genes to Cells 11, (2006).

46. Hickmott, J. W., Gunawardane, U., Jensen, K., Korecki, A. J. & Simpson, E. M. Epistasis between Pax6 Sey and genetic background reinforces the value of defined hybrid mouse models for therapeutic trials. Gene Ther 25, (2018).

47. Paylar, B., Pramanik, S., Bezabhe, Y. H. & Olsson, P.-E. Temporal sex specific brain gene expression pattern during early rat embryonic development. Front Cell Dev Biol 12, 1343800 (2024).

48. Yang, F., Babak, T., Shendure, J. & Disteche, C. M. Global survey of escape from X inactivation by RNA-sequencing in mouse. Genome Res 20, (2010).

49. Lopes, A. M., Arnold-Croop, S. E., Amorim, A. & Carrel, L. Clustered transcripts that escape X inactivation at mouse XqD. Mammalian Genome 22, (2011).

50. Berletch, J. B. et al. Escape from X Inactivation Varies in Mouse Tissues. PLoS Genet 11, (2015).

51. Su, C. H., Dhananjaya, D. & Tarn, W. Y. Alternative splicing in neurogenesis and brain development. Front Mol Biosci 5, (2018).

52. Fleischer, T. C., Weaver, C. M., McAfee, K. J., Jennings, J. L. & Link, A. J. Systematic identification and functional screens of uncharacterized proteins associated with eukaryotic ribosomal complexes. Genes Dev 20, (2006).

53. Simonis, N. et al. Empirically controlled mapping of the Caenorhabditis elegans protein-protein interactome network. Nat Methods 6, (2009).

54. Duek, P. & Lane, L. Worming into the Uncharacterized Human Proteome. J Proteome Res 18, (2019).

55. Huilgol, D. et al. Direct and indirect neurogenesis generate a mosaic of distinct glutamatergic projection neuron types in cerebral cortex. Neuron 111, (2023).

56. Walcher, T. et al. Functional dissection of the paired domain of Pax6 reveals molecular mechanisms of coordinating neurogenesis and proliferation. Development (Cambridge) 140, (2013).

57. Achiro, J. M. et al. Aging differentially alters the transcriptome and landscape of chromatin accessibility in the male and female mouse hippocampus. Front Mol Neurosci 17, (2024).

58. Buonocore, F. et al. Transcriptomic sex differences in early human fetal brain development. Commun Biol 8, 664 (2025).

59. Hill, R. E. et al. Mouse Small eye results from mutations in a paired-like homeobox-containing gene. Nature 354, (1991).

60. Chuma, S. & Nakatsuji, N. Autonomous Transition into Meiosis of Mouse Fetal Germ Cells in Vitro and Its Inhibition by gp130-Mediated Signaling. Dev Biol 229, 468–479 (2001).

61. Collinson, J. M., Hill, R. E. & West, J. D. Different roles for Pax6 in the optic vesicle and facial epithelium mediate early morphogenesis of the murine eye. Development 127, (2000).

62. Rehen, S. K. et al. A new method of embryonic culture for assessing global changes in brain organization. J Neurosci Methods 158, (2006).

63. 63. Andrews S. FastQC A Quality Control tool for High Throughput Sequence Data. https://www.bioinformatics.babraham.ac.uk/projects/fastqc/ (2010).

64. Bolger, A. M., Lohse, M. & Usadel, B. Trimmomatic: a flexible trimmer for Illumina sequence data. Bioinformatics 30, 2114–2120 (2014).

65. Wang, L., Wang, S. & Li, W. RSeQC: quality control of RNA-seq experiments. Bioinformatics 28, 2184–2185 (2012).

66. Kim, D., Paggi, J. M., Park, C., Bennett, C. & Salzberg, S. L. Graph-based genome alignment and genotyping with HISAT2 and HISAT-genotype. Nat Biotechnol 37, 907– 915 (2019).

67. Danecek, P. et al. Twelve years of SAMtools and BCFtools. Gigascience 10, giab008 (2021).

68. Liao, Y., Smyth, G. K. & Shi, W. featureCounts: an efficient general purpose program for assigning sequence reads to genomic features. Bioinformatics 30, 923–930 (2014).

69. Chen, Y., Chen, L., Lun, A. T. L., Baldoni, P. L. & Smyth, G. K. edgeR v4: powerful differential analysis of sequencing data with expanded functionality and improved support for small counts and larger datasets. Nucleic Acids Res 53, gkaf018 (2025).

70. Ge, S. X., Jung, D., Jung, D. & Yao, R. ShinyGO: A graphical gene-set enrichment tool for animals and plants. Bioinformatics 36, (2020).

71. Ochi, S., Imaizumi, Y., Shimojo, H., Miyachi, H. & Kageyama, R. Oscillatory expression of Hes1 regulates cell proliferation and neuronal differentiation in the embryonic brain. Development (Cambridge) 147, (2020).

72. Cheung, T. T., Weston, M. K. & Wilson, M. J. Selection and evaluation of reference genes for analysis of mouse (Mus musculus) sex-dimorphic brain development. PeerJ 2017, (2017).

73. Untergasser, A. et al. Primer3-new capabilities and interfaces. Nucleic Acids Res 40, (2012).

74. Kikkawa, T. et al. Dmrta1 regulates proneural gene expression downstream of Pax6 in the mammalian telencephalon. Genes to Cells 18, (2013).

75. Naher, S. et al. Kinesin-like motor protein KIF23 maintains neural stem and progenitor cell pools in the developing cortex. EMBO J 44, 331–355 (2025).

76. Yamanaka, S. et al. A proximity biotinylation-based approach to identify protein-E3 ligase interactions induced by PROTACs and molecular glues. Nat Commun 13, 183 (2022).

77. Kim, D. I. et al. An improved smaller biotin ligase for BioID proximity labeling. Mol Biol Cell 27, (2016).

78. Wan, L. et al. SRSF6-regulated alternative splicing that promotes tumour progression offers a therapy target for colorectal cancer. Gut 68, (2019).

79. Dong, D. et al. YTHDC1 promotes the malignant progression of gastric cancer by promoting ROD1 translocation to the nucleus. Cell Biol Toxicol 40, 19 (2024).

80. Antona, A. et al. Targeting lysine-specific demethylase 1 (KDM1A/LSD1) impairs colorectal cancer tumorigenesis by affecting cancer cells stemness, motility, and differentiation. Cell Death Discov 9, (2023).

81. Liu, Y. et al. Prognostic potential of PRPF3 in hepatocellular carcinoma. Aging 12, (2020).

82. Xu, N. et al. PUF60 promotes cell cycle and lung cancer progression by regulating alternative splicing of CDC25C. Cell Rep 42, (2023).

83. Tomsic, J. et al. A germline mutation in SRRM2, a splicing factor gene, is implicated in papillary thyroid carcinoma predisposition. Sci Rep 5, (2015).

84. Zhao, Y. et al. The spliceosome factor sart3 regulates hematopoietic stem/progenitor cell development in zebrafish through the p53 pathway. Cell Death Dis 12, (2021).

85. Chang, Y. et al. PHF5A promotes colorectal cancerprogression by alternative splicing of TEAD2. Mol Ther Nucleic Acids 26, (2021).

86. Deka, B. et al. RNPS1 functions as an oncogenic splicing factor in cervical cancer cells. IUBMB Life 75, (2023).

87. Wei, Y., Chen, Z., Li, Y. & Song, K. The splicing factor WBP11 mediates MCM7 intron retention to promote the malignant progression of ovarian cancer. Oncogene 43, 1565– 1578 (2024).

88. Kim, W. et al. PRPF4 Knockdown Suppresses Glioblastoma Progression via the p38 MAPK and ERK Signaling Pathways. Anticancer Res 45, 549–564 (2025).

89. Phillips, J. B. et al. SON is an essential RNA splicing factor promoting ErbB2 and ErbB3 expression in breast cancer. Br J Cancer 131, 1437–1449 (2024).

90. Yang, G.-J. et al. PRMT7 in cancer: Structure, effects, and therapeutic potentials. Eur J Med Chem 283, 117103 (2025).

91. Rengasamy, M. et al. The PRMT5/WDR77 complex regulates alternative splicing through ZNF326 in breast cancer. Nucleic Acids Res 45, (2017).

92. Deshpande, K. et al. SRRM4-mediated REST to REST4 dysregulation promotes tumor growth and neural adaptation in breast cancer leading to brain metastasis. Neuro Oncol 26, (2024).

93. Ma, Z. et al. The role of DDX46 in breast cancer proliferation and invasiveness: A potential therapeutic target. Cell Biol Int 47, (2023).

94. Liu, X. et al. KHDRBS1 regulates the pentose phosphate pathway and malignancy of GBM through SNORD51-mediated polyadenylation of ZBED6 pre-mRNA. Cell Death Dis 15, 802 (2024).

95. Zhi, L. et al. RBM25 depletion suppresses the growth of colon cancer cells through regulating alternative splicing of MNK2. Sci China Life Sci 67, 2186–2197 (2024).

96. Gu, J., Chen, Z., Chen, X. & Wang, Z. Heterogeneous nuclear ribonucleoprotein (hnRNPL) in cancer. Clinica Chimica Acta 507, (2020).

97. Urtasun, R. et al. Splicing regulator SLU7 preserves survival of hepatocellular carcinoma cells and other solid tumors via oncogenic miR-17-92 cluster expression. Oncogene 35, (2016).

98. Zhang, H. et al. DDX39B contributes to the proliferation of colorectal cancer through direct binding to CDK6/CCND1. Cell Death Discov 8, (2022).

99. Zhuang, W. et al. RBM19 promotes the progression of prostate cancer under docetaxel treatment via SNHG21/PIM1 axis. Cell Biol Toxicol 41, 19 (2024).

100. Wang, Z. et al. RBM17 promotes hepatocellular carcinoma progression by regulating lipid metabolism and immune microenvironment: implications for therapeutic targeting. Cell Death Discov 11, 338 (2025).

101. Xu, C. et al. RNA-binding protein 39: a promising therapeutic target for cancer. Cell Death Discov 7, (2021).

102. Thongchot, S., Aksonnam, K., Thuwajit, P., Yenchitsomanus, P. T. & Thuwajit, C. Nucleolin-based targeting strategies in cancer treatment: Focus on cancer immunotherapy (Review). Int J Mol Med 52, (2023).

103. Zhang, Y., Zhang, Z., Dong, J. & Liu, C. HNRNPC as a pan-cancer biomarker and therapeutic target involved in tumor progression and immune regulation. Oncol Res 33, 83 (2024).

104. Tsuboi, M. et al. Ubiquitination-Independent Repression of PRC1 Targets during Neuronal Fate Restriction in the Developing Mouse Neocortex. Dev Cell 47, (2018).

105. Shimojo, H. & Kageyama, R. Real-time Bioluminescence Imaging of Notch Signaling Dynamics during Murine Neurogenesis. JoVE e60455 (2019).

106. Kaya-Okur, H. S. et al. CUT&Tag for efficient epigenomic profiling of small samples and single cells. Nat Commun 10, (2019).

107. Langmead, B., Wilks, C., Antonescu, V. & Charles, R. Scaling read aligners to hundreds of threads on general-purpose processors. Bioinformatics 35, (2019).

108. Dunham, I. et al. An integrated encyclopedia of DNA elements in the human genome. Nature 489, 57–74 (2012).

109. Amemiya, H. M., Kundaje, A. & Boyle, A. P. The ENCODE Blacklist: Identification of Problematic Regions of the Genome. Sci Rep 9, 9354 (2019).

110. Pongor, L. S. et al. BAMscale: quantification of next-generation sequencing peaks and generation of scaled coverage tracks. Epigenetics Chromatin 13, 21 (2020).

111. Robinson, J. T. et al. Integrative genomics viewer. Nat Biotechnol 29, (2011).

