## Supplementary Information for "Sex-biased transcriptome in embryonic mouse cortices under *Pax6* haploinsufficiency highlights Pbdc1 as a candidate regulator"

#### Supplementary Figure 1

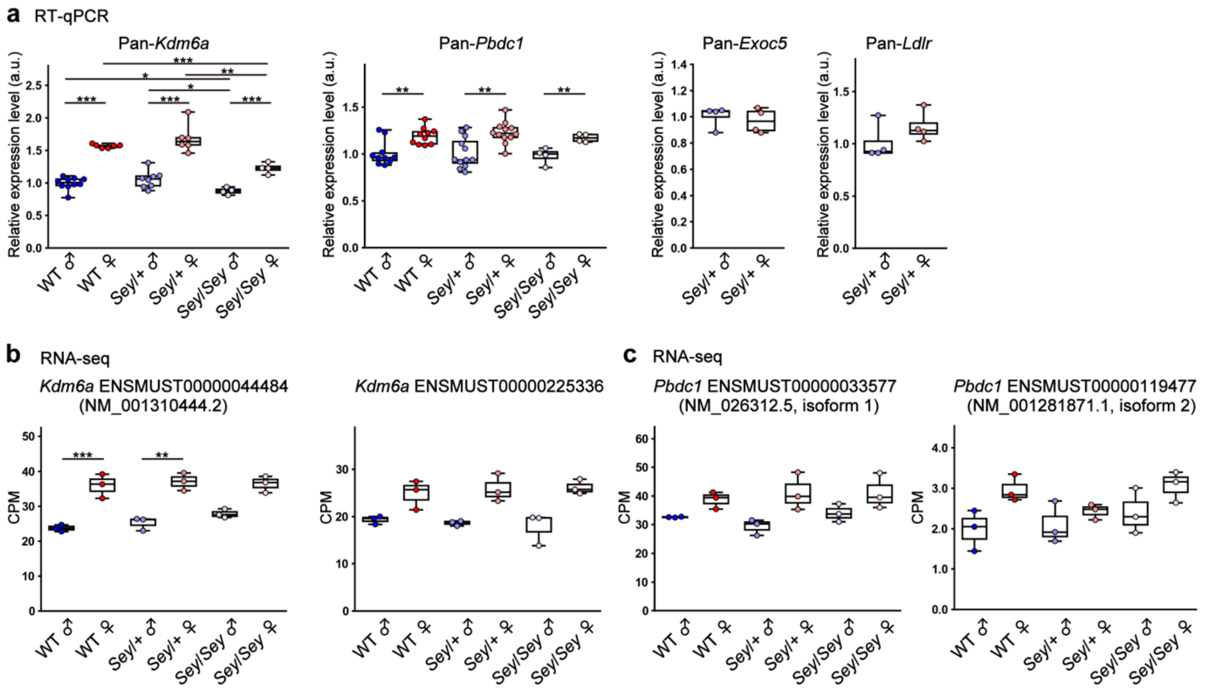


**Supplementary Fig. 1: Expression levels of sex-DEGs in *Sey*/+**

**a** RT-qPCR analyses of pan-*Kdm6a*, pan-*Pbdc1,* pan-*Exoc5 and* pan-*Ldlr* mRNA expression levels in the telencephalons of WT, *Sey*/+ and *Sey*/*Sey*. **b** Isoform-level expression of *Kdm6a* ENSMUST00000044484 (NM_001310444.2), *Kdm6a* ENSMUST00000225336, *Pbdc1* ENSMUST00000033577 (NM_026312.5, isoform 1) and *Pbdc1* ENSMUST00000119477 (NM_001281871.1, isoform 2) in RNA-seq data. $*p < 0.05$, $**p < 0.01$, $***p < 0.001$; determined by two-tailed Student's *t*-test followed by Benjamini-Hochberg multiple testing correction. The line in the middle of box plots represent median, the lower and upper bounds of boxes indicate the first and third quartiles, and end of whiskers denote the minimum and maximum values. CPM, count per million.

#### Supplementary Figure 2

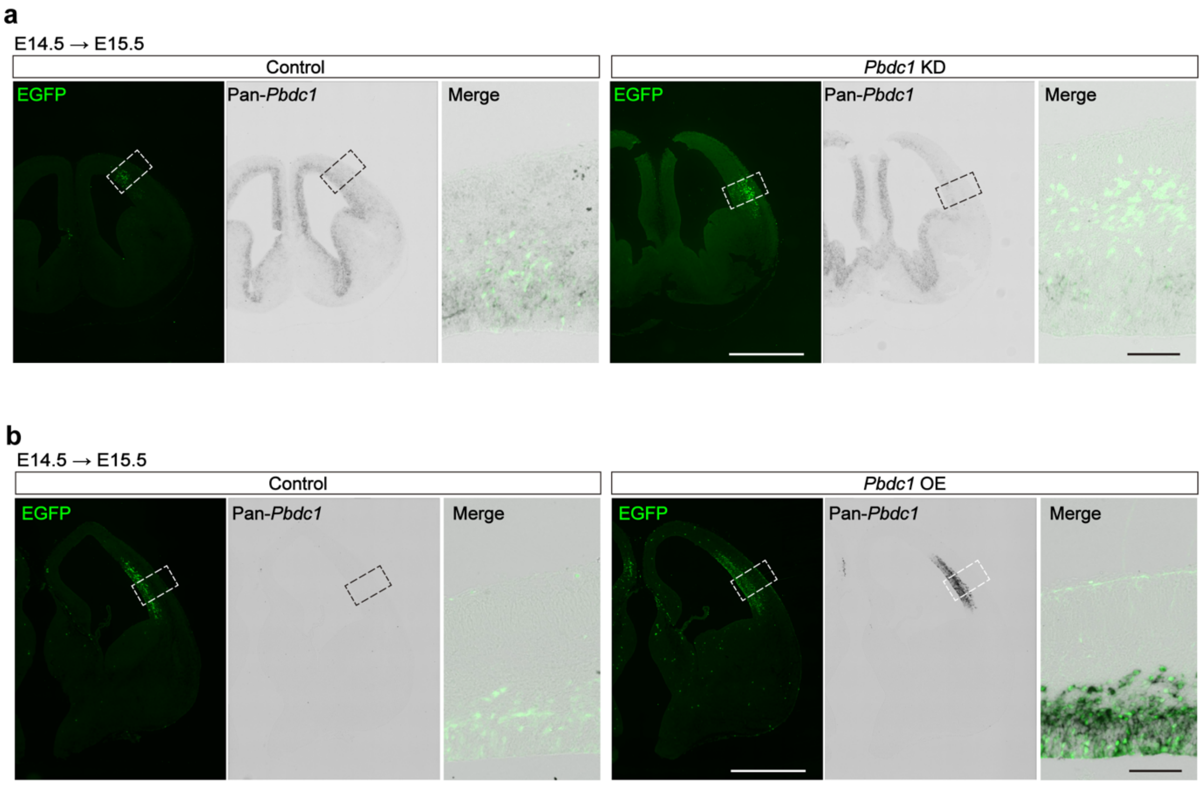


**Supplementary Fig.2: Efficiency of *Pbdc*1-KD and *Pbdc1*-OE at mRNA levels.**

**a** Representative images stained for EGFP protein (green) and pan-*Pbdc1* (black) mRNA in E15.5 cortices at 24 h after *in utero* electroporation of control or *Pbdc1* siRNA. **b** Representative images stained for EGFP protein (green) and pan-*Pbdc1* (black) mRNA in E15.5 cortices at 24 h after *in utero* electroporation of *pCAX* (control) or *pCAX-Pbdc1* vector. Scale bars, lower magnification, 1 mm; higher magnification, 100 μm. Boxes denote the zoomed area.

### Supplementary Tables

**Supplementary Table 1: List of the proteins interacting with Pbdc1 and annotated to RNA-splicing terms.**

**Supplementary Table 2: Count matrix of RNA-seq data at gene-level.**

**Supplementary Table 3: Count matrix of RNA-seq data at isoform-level.**

**Supplementary Table 4: Primer sequences used for RT-qPCR and ChIP-qPCR.**

**Supplementary Table 5: Antibodies used for immunohistochemistry and ChIP.**

**Supplementary Table 6: Quantification of proximity-dependent PPI analysis in N2a cell.**
